# Genomic evolution of antibiotic resistance is contingent on genetic background following a long-term experiment with *Escherichia coli*

**DOI:** 10.1101/2020.08.19.258384

**Authors:** Kyle J. Card, Misty D. Thomas, Joseph L. Graves, Jeffrey E. Barrick, Richard E. Lenski

## Abstract

Antibiotic resistance is a growing health concern. Efforts to control resistance would benefit from an improved ability to forecast when and how it will evolve. Epistatic interactions between mutations can promote divergent evolutionary trajectories, which complicates our ability to predict evolution. We recently showed that differences between genetic backgrounds can lead to idiosyncratic responses in the evolvability of phenotypic resistance, even among closely related *Escherichia coli* strains. In this study, we examined whether a strain’s genetic background also influences the genotypic evolution of resistance. Do lineages founded by different genotypes take parallel or divergent mutational paths to achieve their evolved resistance states? We addressed this question by sequencing the complete genomes of antibiotic-resistant clones that evolved from several different genetic starting points during our earlier experiments. We first validated our statistical approach by quantifying the specificity of genomic evolution with respect to antibiotic treatment. As expected, mutations in particular genes were strongly associated with each drug. Then, we determined that replicate lines evolved from the same founding genotypes had more parallel mutations at the gene level than lines evolved from different founding genotypes, although these effects were more subtle than those showing antibiotic specificity. Taken together with our previous work, we conclude that historical contingency can alter both genotypic and phenotypic pathways to antibiotic resistance.

**Significance:** A fundamental question in evolution is the repeatability of adaptation. Will independently evolving populations respond similarly when facing the same environmental challenge? This question also has important public-health implications related to the growing problem of antibiotic resistance. For example, efforts to control resistance might benefit from accurately predicting mutational paths to resistance. However, this goal is complicated when a lineage’s prior history alters its subsequent evolution. We recently found that differences between genetic backgrounds can lead to unpredictable responses in phenotypic resistance. Here, we report that genetic background can similarly alter genotypic paths to resistance. This historical contingency underscores the importance of accounting for stochasticity, in the past as well as at present, when designing evolutionarily informed treatment strategies.

## Introduction

Convergent evolution is common in nature. The independent emergence of winged flight in insects and mammals, and of camera-like eyes in vertebrates and cephalopod mollusks, are familiar but striking examples of how evolution can drive distantly related lineages to similar phenotypic outcomes (1). For over a century, biologists have sought to understand the processes underlying these patterns and quantify the extent of convergent evolution in the natural world. However, quantifying convergence in nature is difficult for at least two reasons. First, one typically observes only a biased sample of possible outcomes. For example, extinct lineages that evolved different, but ultimately unsuccessful, adaptations usually go undetected, causing one to overestimate the extent of convergence (2, 3). Second, comparative studies generally cannot account for slight differences in environments or in lineages’ prior evolutionary histories as causes of divergent adaptation, leading to an underestimation of convergence (2–4).

Controlled and replicated evolution experiments with microbes allow one to overcome these limitations and more precisely quantify the extent of parallel and convergent evolution (5). Populations of bacteria, yeast, and viruses reproduce quickly and grow to large numbers, permitting one to observe evolution over timescales of days to years. One can also propagate independent lines under identical conditions and characterize evolutionary repeatability at both the phenotypic and genotypic levels.

Evolution experiments with bacteria can also shed new light on the growing crisis of antibiotic resistance. It has been estimated that at least 700,000 people die each year from resistant infections, and the mortality rate has been projected to rise to 10 million by 2050, which would exceed deaths from cancer (6). Alternative strategies are necessary to combat drug resistance, especially given the slowing pace at which new drug classes are being developed. Drug discovery through large-scale chemical screens or isolation of natural products is one approach to addressing the problem of resistance (7). However, bacteria will likely evolve resistance to new antibiotics, as they have to previous ones, limiting the effectiveness of drug discovery over the long-term.

Alternatively, it might be possible to extend the usefulness of existing therapeutics by channeling evolution toward drug susceptible states with an improved understanding of the factors that shape evolutionary trajectories (8–11). To that end, the phenotypic and genetic repeatability of resistance evolution has motivated several studies (12–19). In particular, two landmark studies evaluated the reproducibility of *Escherichia coli* populations evolving in and adapting to increasing antibiotic concentrations in spatially homogeneous (12) and structured environments (15). They found strong signatures of genomic parallelism; that is, replicate lines tended to evolve high-level resistance through mutations in a limited set of genes. However, the replicate lines in these experiments were all founded from genetically identical cells, so it is unknown whether selection would target the same genes if the founding genetic background had been different.

Epistatic interactions between mutations (including between resistance mutations and the genetic backgrounds in which they occur) can affect adaptive trajectories. Thus, they complicate our ability to predict resistance evolution and design effective treatment strategies. In theory, one can leverage collateral drug responses—where the evolution of resistance to one drug increases a bacterium’s susceptibility to other drugs (13, 14, 16, 20–22)—to forestall or even reverse antibiotic resistance in pathogens. Yet differences in genetic background can make this approach difficult in practice. For example, replicate *E. coli* and *Enterococcus faecalis* populations can take different mutational paths to increased resistance that change their collateral responses to second-line antibiotics (18, 19). Despite stochasticity at the level of individual strains, one can still exploit statistical patterns in resistance profiles across many replicates to optimize drug-cycling protocols (18). Recent work showed that such an approach could drive populations to eventual long-term susceptibility through intermediate states of high-level resistance (18).

Replicate populations also accumulate genetic differences as they evolve in permissive environments, and these differences can affect their ability to adapt when challenged with antibiotics. We recently used several strains from the *E. coli* long-term evolution experiment (LTEE) to examine the role that genetic background plays in the evolution of drug resistance (17). In the LTEE, 12 replicate populations were founded from a common ancestral strain and have been propagated daily for over 30 years in an environment without antibiotics (23, 24). Clones from several populations evolved increased antibiotic sensitivity compared to their ancestor (17, 25). We asked whether these strains could compensate for their increased sensitivity through subsequent evolution under drug selection. We found that their evolutionary potential was idiosyncratic with respect to their initial genotype, such that resistance was constrained in some backgrounds but not in others, indicating the role of historical contingency in this process (5, 17).

In this study, we sequenced the complete genomes of antibiotic-resistant clones that evolved from several different founding strains during our earlier experiments and used this information to examine how genetic background affects the genomic evolution of antibiotic resistance. First, we validated our statistical approach by demonstrating that mutations in particular target genes were associated with each of the four antibiotic treatments. We then showed that evolution was also contingent, albeit more subtly, on differences in genetic background, such that resistant lines that evolved from the same genotype had, on average, slightly more mutational targets in common. These results, taken together with our previous work, indicate that even slight differences in genetic background complicate one’s ability to predict phenotypic and genotypic outcomes of antibiotic resistance evolution.

## Results

### Genomic Evolution of Strains Evolved under Four Different Antibiotic Treatments

In our previous study (17), we isolated antibiotic-resistant mutants that evolved from five different parental genotypes: the LTEE ancestor and four derived clones isolated from the LTEE at generation 50,000. The experiment was performed over one round of drug selection. Here, we sequenced the complete genomes of 64 of the resistant clones, but we discarded 3 that were identified as cross-contaminants (Materials and Methods) (Table S1).

The 61 remaining resistant clones had a total of 76 mutations. Forty-five genomes had a single mutation, 11 others had two mutations, and 3 genomes had three mutations (Fig. 1). Two other clones (both in the tetracycline treatment) had no identifiable mutations; they might have had unstable genetic changes or types of mutations that could not be resolved by our analyses of the short-read sequencing data (see Materials and Methods). In any case, we have excluded these two clones from the analyses that follow. Twenty-seven of the 76 mutations (35.5%) were single-base substitutions; of these, 22 were either nonsynonymous or nonsense mutations that altered the encoded protein’s amino-acid sequence, 1 was synonymous, and 4 occurred in *alaT*, which encodes a tRNA rather than a protein. Five other mutations (6.6%) were in intergenic regions within 150-bp upstream of a gene, which suggests that they affect regulation.

**Fig. 1.**
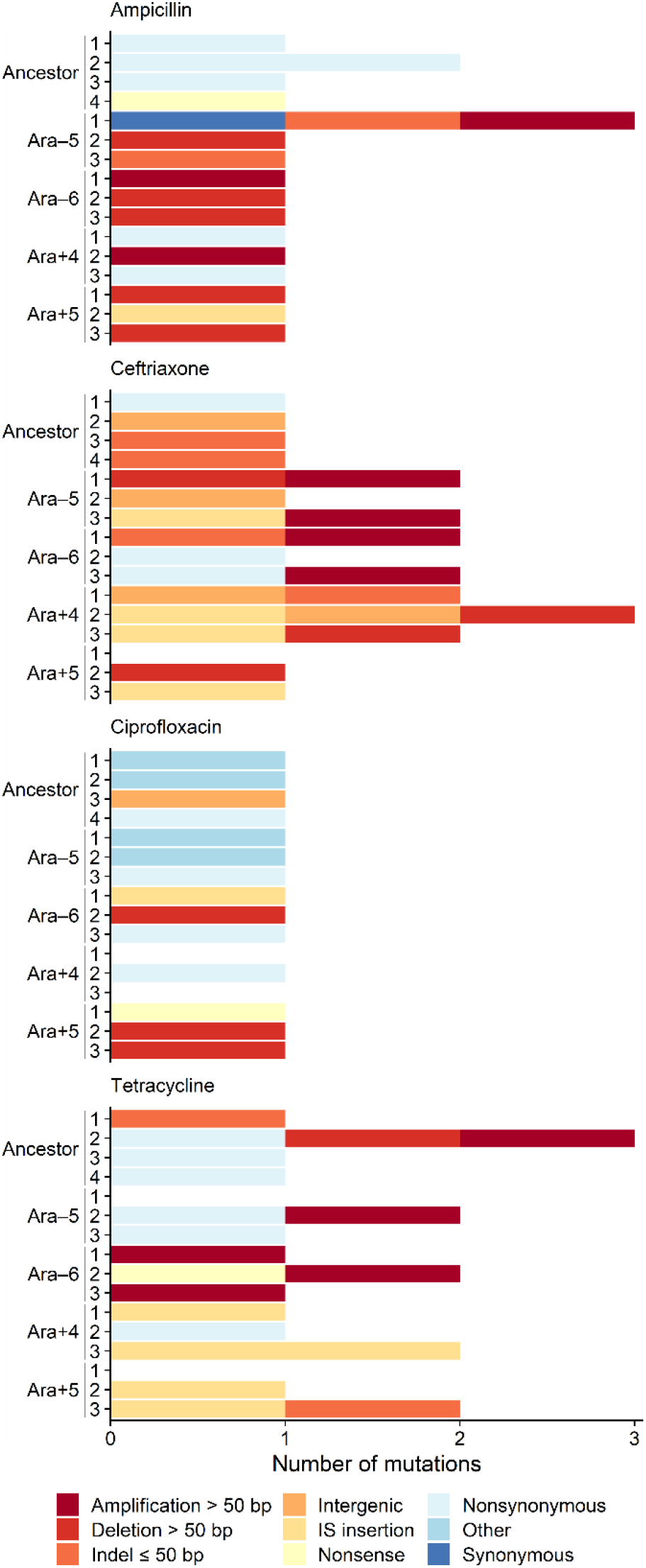
Numbers and types of mutations in evolved genomes. Summary of the 76 mutations observed in 61 antibiotic-resistant clones after selection in ampicillin, ceftriaxone, ciprofloxacin, or tetracycline. Mutations are color-coded by the type of genetic change, according to the legend at the bottom. The “Other” category represents mutations in a tRNA. Evolved genomes are labeled according to their parental genetic background and replicate. Two tetracycline-selected clones (Ara–5-1 and Ara+5-1) had no identifiable mutations (see Materials and Methods).

The largest proportion of mutations in the resistant lines were structural variants. They comprise 44 (57.9%) of the observed changes (Fig. 1). Eleven of these were IS-element insertions in protein-coding genes; 8 were small insertions and deletions (indels) of less than 50 bp, 13 were large deletions, and 12 were large amplifications. Twelve of the 13 large deletions and 11 of the 12 large amplifications were found in lines derived from the generation-50,000 backgrounds (Fig. 1). However, 45 of the 61 clones (73.8%) belong to that group, and neither observed distribution deviates significantly from that null expectation (binomial tests, *p* = 0.1077 and *p* = 0.1368 for large deletions and large amplifications, respectively).

### Genomic Parallelism at the Functional Level

Antibiotic resistance can arise through mutations that change gene regulation and expression, cell permeability and efflux, and metabolism (26). To determine how drug selection acted on these functions in our experiment, we quantified the extent of genomic parallelism in the resistant lines at the functional level (i.e., sets of genes that share broadly defined functions). We used the curated descriptions of cellular processes in EcoCyc (27) to match each mutated gene to an associated function. We excluded large deletions and amplifications when the affected genes do not share a common function.

About 37% of the 57 mutations that fit the criteria for inclusion occurred in regulatory genes, ~26% in metabolic genes, ~21% in genes that encode transporters, and ~11% in genes involved in transcription or translation (Fig. 2). More mutations in some of these functional categories than in others might suggest a pattern of parallel evolution. However, more *E. coli* genes are involved in some functions than in others, and therefore a random mutation is more likely to occur in those categories that constitute a larger proportion of the genome. To examine whether the observed number of mutations in each category occurred more frequently than expected (28), we modeled the data using the Poisson cumulative expectation *P*(*x ≥ observed, λ*), where *λ* is given by: (total number of mutations) × [(combined length of all genes in functional category) / (total genome length)]. The derived parental backgrounds evolved slightly smaller genomes during the LTEE (24), so we used their average genome size (4,602,572 bp) when calculating *λ*.

**Fig. 2.**
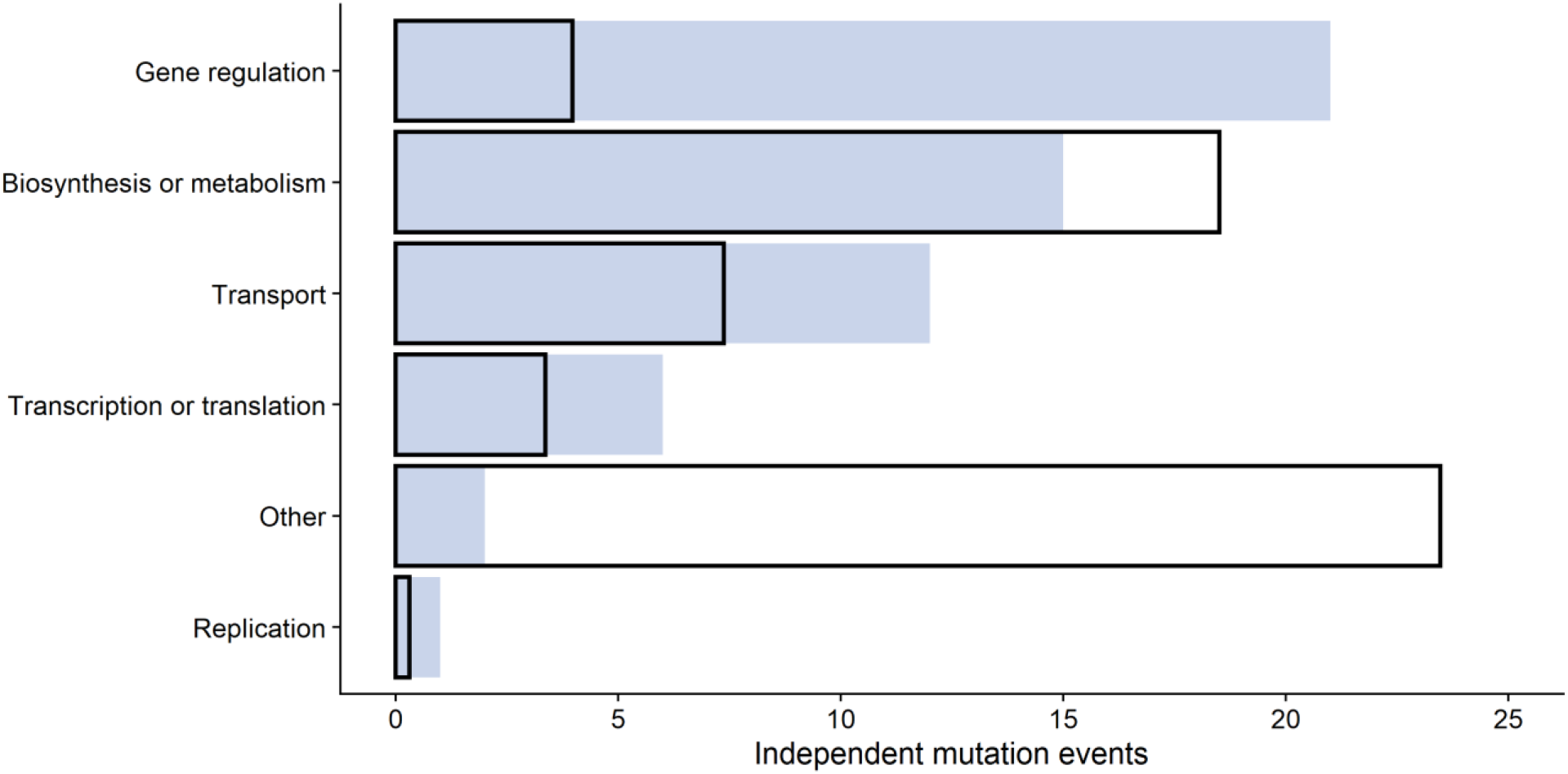
Distribution of independent mutations in different functional categories. The observed and expected distributions are shown as shaded regions and outlines, respectively. See text for statistical analysis.

Regulatory genes accrued mutations about 5 times more often than expected from the Poisson distribution (Fig. 2), and this difference is highly significant (*p* < 0.0001). Genes involved in transport functions had about 1.6 times more mutations than expected by chance, but this difference was marginally non-significant (*p* = 0.0723). Genes involved in transcription or translation had about 1.8 times as many mutations as expected, but this difference was also not statistically significant given the small number of mutations in these targets (*p* = 0.1245). It should be emphasized that this analysis is conservative because it lumps together all of the genes in each functional category. However, mutations in only a subset of these genes are likely to cause resistance. Therefore, the effective mutational target size and the resulting expected number of mutations is presumably much smaller.

### Specificity of Genomic Evolution in the Different Antibiotic Environments

We compared the gene-level similarity of mutations between independent lines that evolved in the same antibiotic treatment and across the four different treatments to evaluate the effect of the selective environment on the genetic paths to increased antibiotic resistance. As described in the Materials and Methods, we computed Dice’s coefficient of similarity for each pair of clones using the 71 qualifying mutations that could be assigned to a particular gene. The average within-treatment similarity was 0.089 and the average between-treatment similarity was 0.032 (Fig. 3). In other words, two clones that independently evolved under the same antibiotic selection had on average 8.9% of their mutated genes in common, whereas those that evolved under different antibiotics shared on average only 3.2% of their mutated genes. A randomization test shows that the 5.7% difference in similarity is highly significant (*p* < 0.0001). Thus, genomic evolution was demonstrably specific with respect to the antibiotic treatment.

**Fig. 3.**
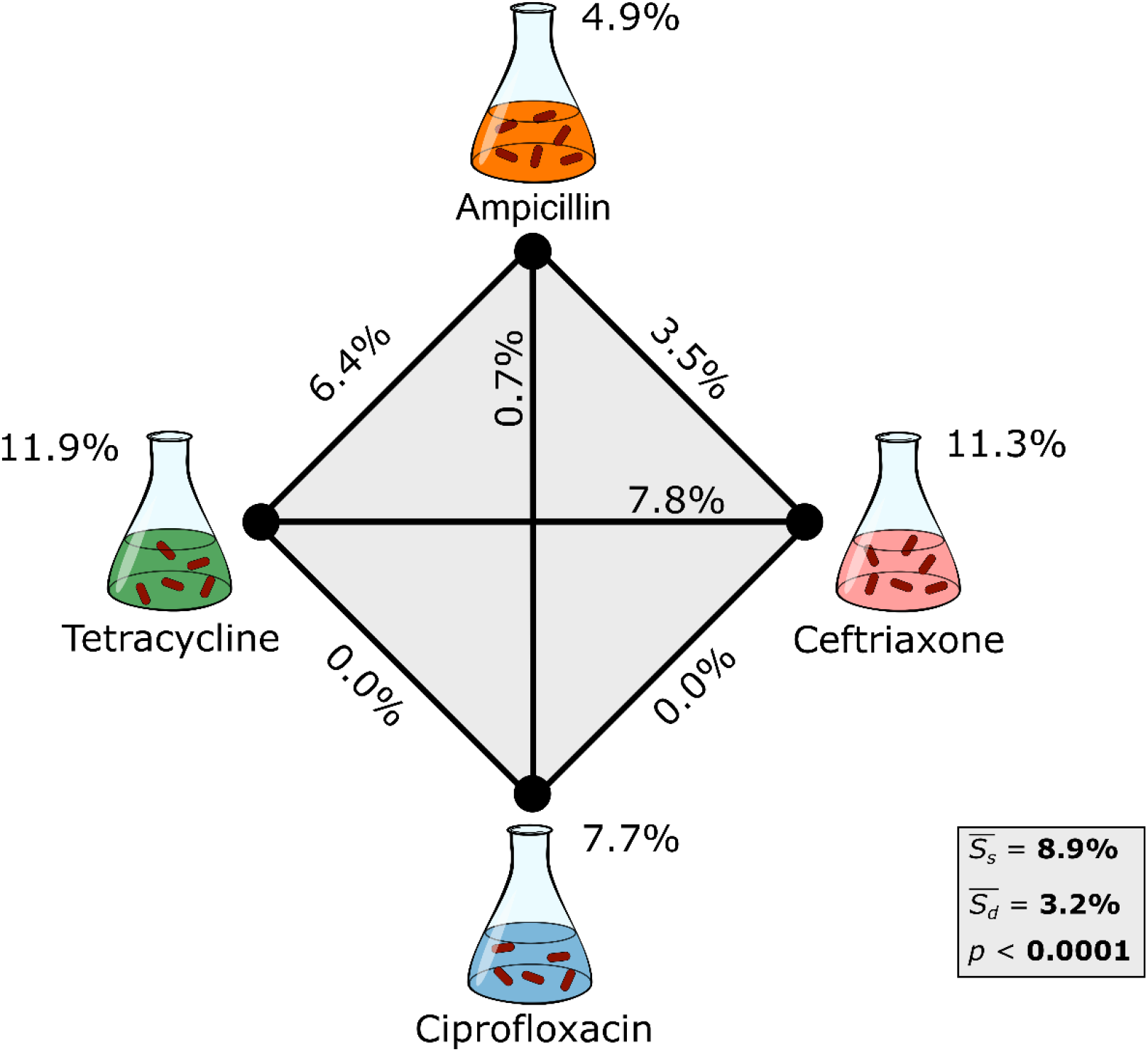
Specificity of genomic evolution with respect to antibiotic treatment. Treatments and the edges connecting them are labeled with Dice’s coefficient scores that show the average similarity for all clone pairs evolved in the same antibiotic (*S_s_*) and in different antibiotics (*S_d_*), respectively. Only the 71 qualifying mutations (see Materials and Methods) were included in the calculations. The weighted averages of *S_s_* and *S_d_* are shown in the grey box. The difference between these two values indicates the extent to which genome evolution was specific to the antibiotic treatment. The resulting *p*-value was calculated using a randomization test.

The similarity analysis does not reveal the specific genes that contribute to the antibiotic-treatment specificity. To address this issue, we used Fisher’s exact tests to identify genes that had an excess of qualifying mutations in the replicate lines evolved under the four treatments (Fig. 4). We found 5 “signature” genes in which mutations contributed significantly to antibiotic specificity (Table 1, Fig. 4). The *alaT* gene encodes a tRNA; it was mutated in 4 of the 14 CIP-resistant lines, but in none of the other 44 lines with qualifying mutations (Fig. 4). The *ompR* gene is part of the two-component system that regulates the production of outer-membrane proteins; it was mutated in 6/14 TET-resistant lines as well as in 4/44 lines that evolved resistance to other drugs. The other gene in this two-component system, *envZ*, was mutated in 2 of the 10 TET-resistant lines that did not have an *ompR* mutation. Two genes, *ompF* and *hns*, were associated with resistance to ceftriaxone (Table 1). The former encodes an outer-membrane porin and was mutated in 6/15 CRO-resistant lines along with 3/43 other lines; the latter encodes a histone-like global regulator and acquired mutations in 3/15 CRO-resistant lines and 1 of the 43 lines that became resistant to another antibiotic (Fig. 4). Finally, a large deletion was found in 3 of the 15 AMP-resistant lines but not in any of the other 43 lines (Table 1); this deletion affects multiple genes including *phoE*, which encodes an outer membrane porin (Fig. 4).

**Table 1.**
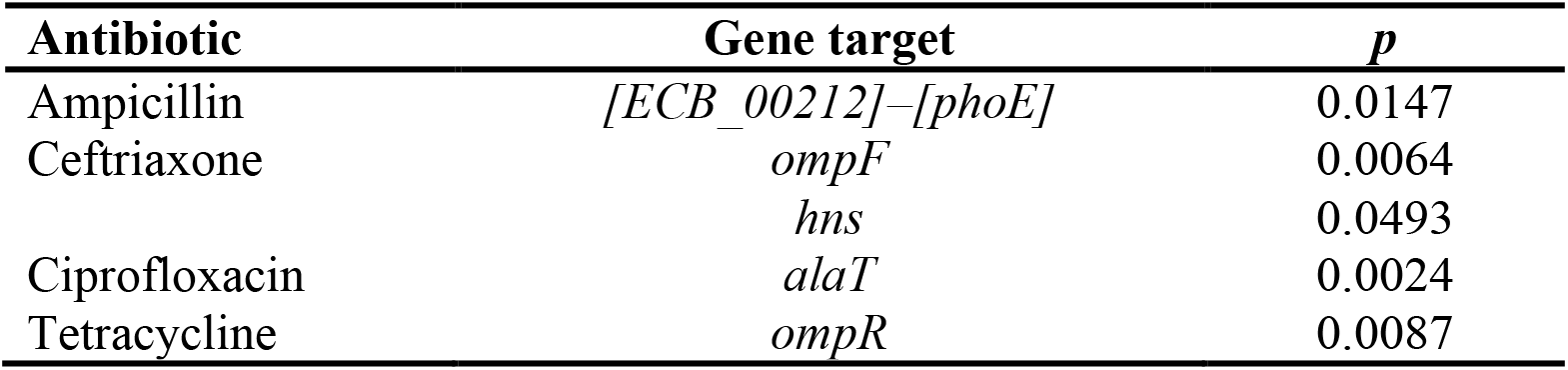
Gene targets that contributed to antibiotic-treatment specificity. Statistical significance was calculated using Fisher’s Exact Test for the association between antibiotic treatments and gene targets.

| Antibiotic | Gene target | *p* |
| --- | --- | --- |
| Ampicillin | [*ECB_00212*]-[*phoE*] | 0.0147 |
| Ceftriaxone | *ompF* | 0.0064 |
|  | *hns* | 0.0493 |
| Ciprofloxacin | *alaT* | 0.0024 |
| Tetracycline | *ompR* | 0.0087 |

**Fig. 4.**
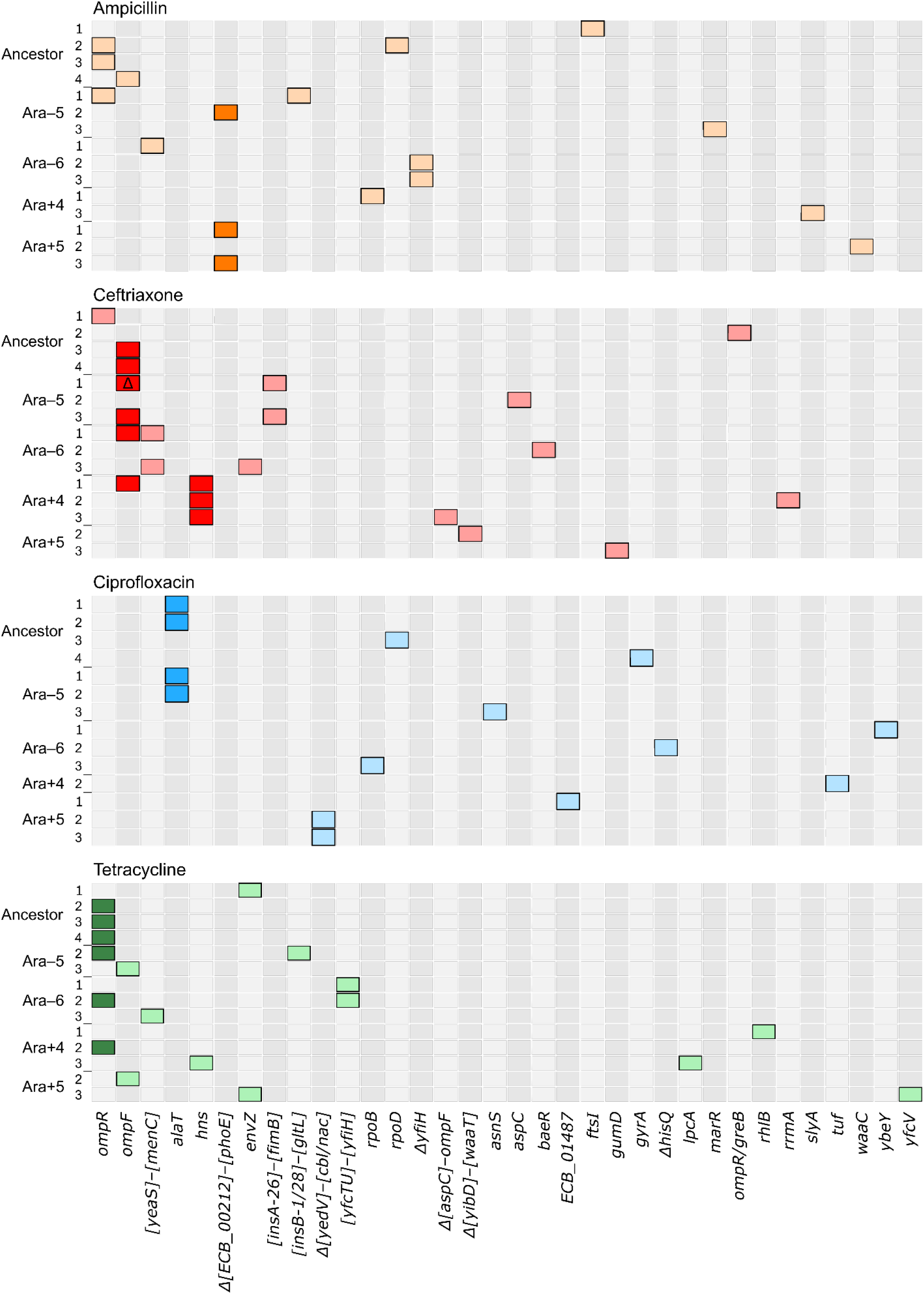
Identity of mutated genes in antibiotic-resistant lines. A total of 58 lines (labels at left) evolved from 5 different genetic backgrounds in ampicillin, ceftriaxone, ciprofloxacin, or tetracycline environments. Two (TET Ara–5-1, TET Ara+5-1) had no identifiable mutations; a third (AMP Ara+4-2) had no qualifying mutation that could be assigned to a specific gene (see Materials and Methods). These three lines are not shown. Filled cells identify the 71 qualifying mutations by the affected genes (shown along the bottom and listed in order of the total number of mutations). The darkly shaded cells identify signature genes, in which mutations are significantly associated with one antibiotic treatment (Table 1). A deletion or amplification spanning a given genomic region is indicated when two gene names are shown. If a gene name is shown in brackets, then only part of that gene is affected. If a gene name is preceded by Δ, then those genes are deleted; otherwise, they are amplified. Part of the *ompF* gene is deleted in the CRO Ara–5-1 line.

### Specificity of Genomic Evolution with Respect to Genetic Background

We next employed Dice’s coefficient of pairwise similarity to quantify the specificity of genomic evolution with respect to the parental strain’s genetic background. Using the ampicillin treatment as an example (Fig. 5A), the clones that evolved independently from the same founding genetic background and from different backgrounds had, on average, 14.8% and 3.1% of their mutated genes in common, respectively, indicating a difference of 11.7%. This trend of greater similarity for clones derived from the same genetic background also occurred in the three other antibiotics (Fig. 5B–D).

**Fig. 5.**
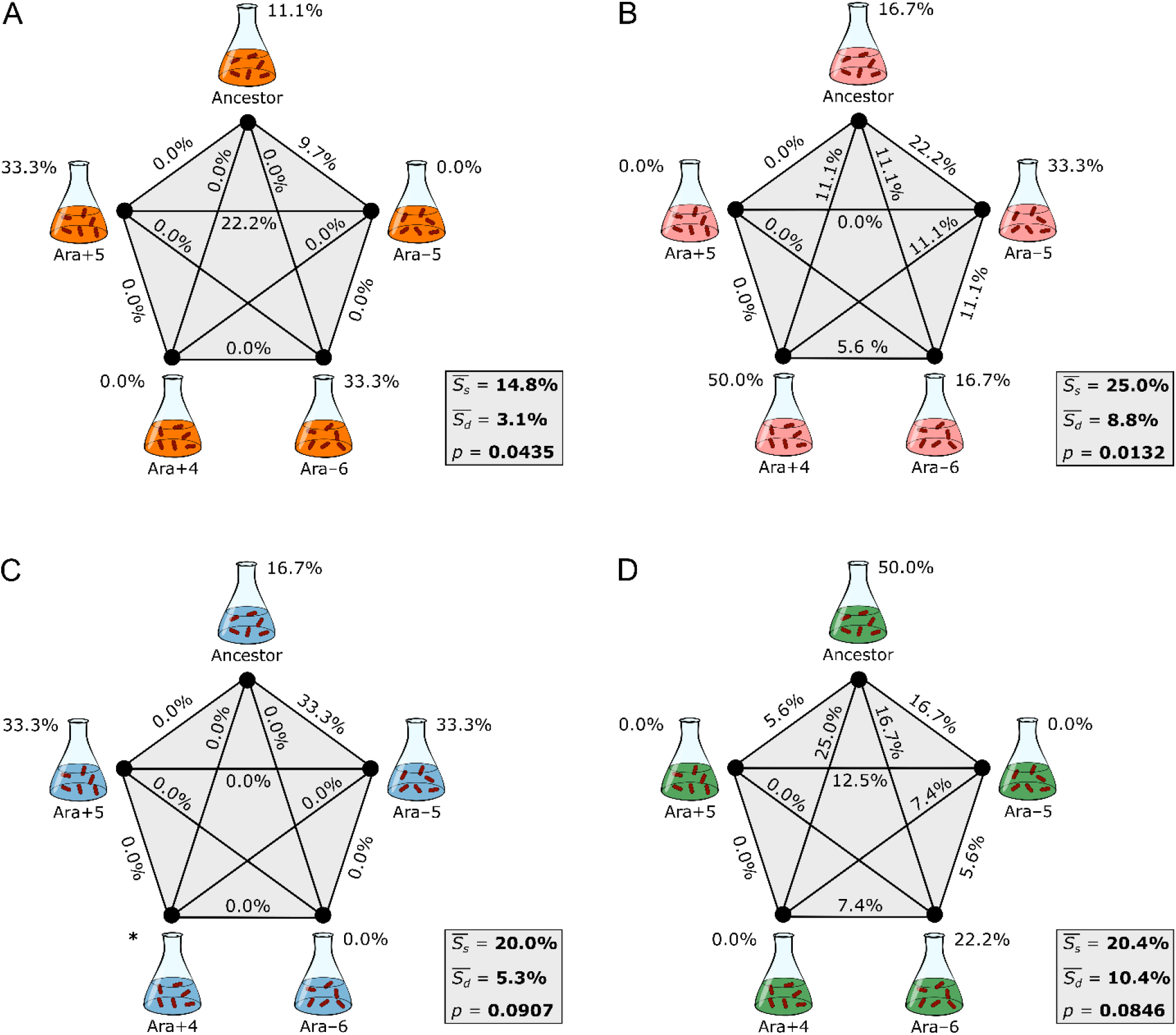
Specificity of genomic evolution with respect to genetic background. Five different backgrounds and the edges connecting them are labeled with Dice’s coefficient scores that show the average similarity for all clone pairs evolved from the same genetic background (*S_s_*) and from different backgrounds (*S_d_*), respectively, in ampicillin (A), ceftriaxone (B), ciprofloxacin (C), and tetracycline (D). The difference between *S_s_* and *S_d_* indicates the extent to which genome evolution was specific to the genetic background. Two of the three replicates derived from the Ara+4 background in ciprofloxacin were excluded owing to cross-contamination, and *S_s_* cannot be calculated in that case (*). See Fig. 3 for additional details.

We performed separate randomization tests for the clones in each antibiotic treatment to evaluate whether the effects of genetic background were significant. The associations between genomic evolution and the identity of the parental strain were significant for the lines that evolved in the ampicillin and ceftriaxone environments (Fig. 5A and 5B), and they were marginally non-significant for the lines in the ciprofloxacin and tetracycline environments (Fig. 5C and 5D). When we combined the probabilities from these four independent tests of the hypothesis that differences in genetic background influence the genetic basis of antibiotic resistance using Fisher’s method (29, 30), the overall trend toward greater similarity (gene-level parallelism) of lines evolved from the same founding genotype was highly significant (χ^2^ = 24.67, df = 8, *p* = 0.0018).

In our analysis of genome specificity with respect to antibiotic treatment, we identified several signature genes that contributed significantly to that specificity (Table 1, Fig. 4). We have much less statistical power to identify particular genes that contribute to specificity with respect to genetic background, because only 3 or 4 replicate lines derive from any given background in each antibiotic treatment. Nonetheless, we can identify candidate loci that may contribute to that specificity, which might be further studied in the future. Table 2 shows all of the genes that fulfilled both of the following criteria for a given antibiotic treatment: (i) two or more lines derived from the same background had mutations affecting the same gene; and (ii) that background produced at least as many mutations affecting that gene as did the other four backgrounds combined. In the case of each antibiotic treatment, at least two genetic backgrounds have candidate signature genes that fulfill these criteria.

**Table 2.** Candidate genes that may contribute to genetic-background specificity.

| Antibiotic | Genetic background | Gene target |
| --- | --- | --- |
| Ampicillin | Ancestor | *ompR* |
| Ampicillin | Ara-6 | *yfiH* |
| Ampicillin | Ara+5 | *[ECB_00212]–[phoE]* |
| Ceftriaxone | Ara-5 | *[insA-26]–[fimB]* |
| Ceftriaxone | Ara-6 | *[yeaS]–[menC]* |
| Ceftriaxone | Ara+4 | *hns* |
| Ciprofloxacin | Ancestor | *alaT* |
| Ciprofloxacin | Ara-5 | *alaT* |
| Ciprofloxacin | Ara+5 | *[yedV]–[cbl/nac]* |
| Tetracycline | Ancestor | *ompR* |
| Tetracycline | Ara-6 | *[yfcTU]–[yfiH]* |

## Discussion

How does genetic background affect the evolution of antibiotic resistance? We previously addressed this question by examining the resistance potential of the *E. coli* ancestor of a long-term experiment and derived clones isolated from four populations after generation 50,000. We challenged these strains using a series of drug concentrations, and we found that several strains had a reduced capacity to evolve resistance relative to their ancestor, implicating the role of historical contingency in this process (5, 17, 31). In this study, we asked whether genetic background also influences the genomic basis of resistance by channeling evolution along different mutational paths. We sequenced the complete genomes of 61 resistant lineages that evolved in our earlier experiment to identify the mutations that conferred resistance. We then analyzed whether there were particular signatures of (i) the antibiotic treatment and (ii) the initial background evident from the identities of the mutated genes.

The populations that evolved resistance to the four different drugs in our study exhibited divergent underlying genetic changes (Fig. 3). This result was expected given that bacteria generally evolve resistance through mutations specific to a drug’s mechanism of action (12, 15, 21). This specificity was driven by parallel mutations in several genes (Table 1). Overall, *ompR* and *ompF* had more mutations than any other genes (Fig. 4). OmpR is a transcriptional regulator involved in responses to osmotic and acid stress; mutations to OmpR also contribute to antibiotic resistance by altering the expression of the OmpF major porin (32–34). A recent study showed that *ompF* deletions reduce the permeability of β-lactams (e.g., ampicillin and ceftriaxone) across the outer membrane, thus increasing resistance (34). We found that *ompF* mutations were strongly associated with ceftriaxone-resistant lines, consistent with this prior study. However, the evolution of ampicillin resistance occurred through more diverse mutational paths in our experiment. Although there were some mutations in *ompF* and *ompR* in the ampicillin treatment, we saw a significant association of that treatment with deletion of a different outer-membrane porin, PhoE. The down-regulation of this porin also partially modulates the cell’s response to osmotic stress (35). In the tetracycline treatment, mutations were more common in *ompR* than in *ompF*, which suggests that altering the expression of other genes in this regulon also contributes to resistance. Mutations in *hns* were associated with ceftriaxone-resistant lines. Nishino and Yamaguchi (36) showed that deletion of this global transcriptional regulator increases resistance to multiple drugs because it causes overexpression of several efflux pumps. In contrast to the results for the other three antibiotics in our study, the evolution of ciprofloxacin resistance was not associated with mutations in genes related to outer-membrane proteins. Instead, mutations in *alaT*, which encodes an alanine tRNA, were a signature of this treatment. The mechanism behind this resistance is unknown. One possibility is that these mutations modulate interactions that have been reported between this tRNA and tmRNA, which rescues stalled ribosomes from aberrant translational events (37). Enhanced rescue might directly promote survival or indirectly affect the expression of other vital genes, when cells are treated with this antibiotic.

The signature genes that we observed for each treatment are not the canonical resistance genes for the respective antibiotics (26). Ampicillin and ceftriaxone irreversibly bind to transpeptidases and disrupt cell-wall synthesis; ciprofloxacin targets topoisomerase and inhibits DNA replication; and tetracycline targets the ribosome and hinders protein synthesis. Drug resistance often arises through modifications to these targets, yet these changes rarely occurred in our study. This discrepancy may reflect two factors, one environmental and the other genetic. First, altering a drug target often confers high-level resistance, but at the expense of bacterial growth rate (10, 38). We used moderate drug concentrations to select for mutants (17), and the observed resistance rarely reached levels defined as clinically relevant (39). This moderate environment should favor mutations that provide sufficient resistance at a low fitness cost, because they will leave more descendants during population growth before treatment, and consequently they will be seen more often after the antibiotic challenge. Second, the *E. coli* used in our experiments are all derived from a B strain that differs in important ways from the K-12 strains that are more widely used in studies of antibiotic resistance (40). In particular, *E. coli* K-12 has two major porins, OmpC and OmpF, whereas *E. coli* B expresses only OmpF (41, 42). Thus, the use of the *E. coli* B strain background may well have influenced which genes could mutate to yield resistance in our experiments.

We also found genomic signatures of adaptive divergence associated with differences in genetic background, and these differences are far smaller than those between *E. coli* B and K-12. We sequenced and analyzed resistant lines that evolved from five backgrounds that were separated in time by only a few decades, and which differed only in the mutations that had accumulated in the antibiotic-free environment of the LTEE (Fig. 1). Three or four resistant lines independently evolved from each parental background for each of the four antibiotics studied, allowing us to assess the genomic specificity of resistance with respect to the genetic background (Fig. 5). Although these background effects were more subtle than those showing antibiotic specificity, they are compelling when taken together. Various factors might contribute to the genetic background specificity. Most broadly, epistatic interactions can cause the same mutation to have different effects on resistance, or on its fitness costs, in different backgrounds. The rates at which particular resistance mutations arise may also vary between different genetic backgrounds.

Imagine that the same mutation arises in separate populations founded from two distinct backgrounds. If the mutation confers less resistance in one background than the other, then it may go undetected when those populations are challenged at a high drug concentration. This type of epistasis could therefore generate a signature of genomic specificity of resistance mutations with respect to the genetic background. It is also possible that different genetic backgrounds affect the evolution of resistance by changing the likelihood of certain genomic amplifications or deletions. These types of structural mutations often occur by homologous recombination between IS elements, and they can confer resistance by altering the number of membrane transporters or drug targets (43). Such mutations can also occur spontaneously at very high rates in comparison to point mutations (43, 44). In our study, many of the resistance mutations were mediated by IS elements, including new copies in the derived backgrounds that previously arose during the LTEE (24). The evolution of resistance in these cases is therefore influenced, at least in part, by changes in the rates at which certain types of mutations arise in the derived genetic backgrounds.

Antibiotic resistance is a growing public-health concern. If the most likely evolutionary paths to resistance can be accurately predicted, then there exists a potential opportunity to control the emergence of resistance through rational treatment strategies. However, to predict the evolution of resistance with accuracy, we must understand and integrate information about many factors, including a bacterial lineage’s evolutionary history, and how that history may potentiate or constrain its future evolution. The results from this study, together with our previous findings, demonstrate the importance of historical contingency in the evolution of drug resistance at both the phenotypic and genotypic levels. This contingency underscores the importance of accounting for stochasticity, in the past as well as at present, when designing evolutionarily informed treatment strategies.

## Materials and Methods

### Evolution Experiments and Bacterial Strains

The LTEE has been described in detail elsewhere (3, 23). Briefly, 12 replicate populations of *E. coli* were founded from a common ancestral strain called REL606. These populations have been propagated for more than 70,000 bacterial generations by daily 1:100 transfers in a glucose-limited Davis minimal (DM) medium without antibiotics.

In a previous study (17), we measured the intrinsic resistance of the LTEE ancestor and derived clones isolated from four populations (designated Ara–5, Ara–6, Ara+4, and Ara+5) at generation 50,000 to the antibiotics ampicillin (AMP), ceftriaxone (CRO), ciprofloxacin (CIP), and tetracycline (TET). We also quantified these strains’ capacities for evolving resistance by challenging them across a range of concentrations to these same drugs during one round of selection. In this study, we sequenced the complete genomes of a subset of the resistant mutants that evolved during these experiments, and we examined whether the genetic targets of the resistance mutations systematically differed between the four antibiotics and five genetic backgrounds. Specifically, for each antibiotic treatment we sequenced 4 mutants that independently evolved from the ancestral background, and 3 mutants from each derived background, for a grand total of 64 sequenced mutants (16 mutants × 4 antibiotics) (Table S1).

### Library Preparation and Genome Sequencing

Glycerol stocks of frozen samples were grown overnight in 3 mL of Luria Bertani (LB) medium at 37°C with shaking at 250 rpm. Overnight cultures were harvested by centrifugation and genomic DNA was extracted using the E.Z.N.A. Bacterial DNA kit (Omega Bio-tek). DNA was quantified using the QuantiFluor dsDNA system (Promega). Then, 250 ng of purified genomic DNA was used from each sample for library preparation using the Nextera DNA Flex Library Prep Kit (Illumina) per the manufacturer’s protocols. The 12 pM final libraries were loaded into a 600-cycle V3 MiSeq reagent cartridge and sequenced on an Illumina MiSeq at North Carolina A&T State University. The resultant FASTQ files of 300-base paired-end reads were deposited to the NCBI Sequence Read Archive (accession number: PRJNA649277).

### Mutation Identification

We trimmed sequencing reads to remove low-quality bases using Trimmomatic v0.38 (45) on the Galaxy web platform (46). A sliding-window approach was used; reads were clipped when the average quality score was < 20 in a 4-bp window and to a minimum length of 36 bp. Mutations were then identified in the genomes using *breseq* v0.35 (47) with default parameters. We used a version of the REL606 reference genome with updated gene annotations for variant calling (24, 48).

Each population evolved unique substitutions in *pykF* during the LTEE that distinguish them from one another (24). Therefore, we first compared this locus for each resistant clone against its corresponding parental strain to test for possible external and cross-contamination. Strains KJC184 and KJC217 from the CIP and CRO treatments, respectively, were supposed to derive from the Ara+4 and Ara+5 parental backgrounds, respectively, but they had *pykF* alleles corresponding to other backgrounds used in this study. Also, strain KJC152 from the CIP treatment was supposed to derive from the Ara+4 background, but its genome was identical to a resistant mutant derived from the ancestral clone. We discarded these three cross-contaminants from our study.

The *breseq* results for each of the other 61 sequenced resistant clones gave information on both its genetic background and the mutations that evolved during our previous antibiotic-selection experiments. We manually curated the results by removing the background-specific mutations (i.e., those that arose during the LTEE), which we did by comparing each resistant clone to its parental strain. We also excluded expansions and contractions of hypermutable short sequence repeats that are unlikely to contribute to stably inherited resistance, and mutations within multi-copy elements (e.g., ribosomal RNA operons and insertion sequences) that may result from gene conversions but cannot be fully resolved using short-read sequencing data. In addition, we resolved numerous structural variants by manually examining the depth of read coverage across the genome and predictions of new sequence junctions from split-read mapping for each clone (47). To verify the predicted mutations, we applied the genomic changes in each parental background to the REL606 reference genome and reran *breseq*.

In total, we identified mutations in 59 of the 61 antibiotic-resistant clones. Two clones (KJC65 and KJC66) had no clear genetic changes despite exhibiting increased phenotypic resistance in our earlier study (17). This discrepancy suggests at least two possible explanations. First, these resistant lines might have mutations that could not be adequately resolved by short-read sequencing, including amplifications of genes or chromosomal regions and inversions bounded by identical sequences (e.g., multi-copy IS elements) (49). Second, these lines might have had unstable genetic changes, including copy number changes in homopolymeric tracts and gene amplifications, which are often unstable and can lead to hypermutability, phase variation, and other complications (43, 50–52). To look for amplifications that might have been missed by the *breseq* pipeline, we also used another pipeline (53) to examine the two genomes without identifiable mutations for evidence of regions with above-average read coverage. However, this analysis did not reveal any amplifications in these two clones. Additional details can be found in the R Notebook provided on GitHub (https://github.com/KyleCard/LTEE-ABR-mutant-sequencing).

### Statistical Methods

We quantified the specificity of genomic evolution with respect to antibiotic treatment by comparing the gene-level similarity of mutations between independent resistant lines that evolved under the same treatment versus different treatments. Similarly, for each antibiotic we quantified the specificity of genomic evolution with respect to genetic background by comparing the gene-level similarity of mutations between lines derived from the same parental genotype versus different parental genotypes. Following Deatherage et al. (54), we included in our comparisons nonsynonymous point mutations, small indels, and IS element insertions in genes or within 150-bp upstream of the start of a gene. However, we modified their approach to also include large deletions and amplifications if at least one of the affected genes was also found to be mutated in another clone or if there were parallel changes across lines. We excluded from these analyses synonymous mutations, the two clones with no identified genetic changes, and a third clone with only a large amplification that was unique and could not be assigned to any particular gene. A total of 71 mutations qualified based on these criteria.

We then calculated Dice’s coefficient of similarity, *S*, for each pair of evolved clones, where *S* = 2|*X* ∩ *Y*|/(|*X*| + |*Y*|). Here, |X| and |Y| represent the number of genes with qualifying mutations in each clone, and |*X* ∩ *Y*| is the number of mutated genes in common between them. *S* therefore ranges from 0, when the pair of clones have no mutated genes in common, to 1, when both have mutations in exactly the same set of genes (30, 53, 54). Finally, we calculated the average of these coefficients for all pairs of clones evolved within the same treatment or from the same parental genotype, *S_s_*, and for all pairs of clones evolved across different treatments or different genotypes, *S_d_*. The difference between these two values serves as a test statistic for the specificity of genomic evolution.

The observed outcome can be seen as one of many possible but equally likely outcomes that could have arisen by chance. One can therefore perform a randomization test to evaluate the significance of the test statistic associated with the observed outcome (30). To do so, we repeatedly rearranged the clones associated with each antibiotic treatment, or the clones within each treatment when testing for background specificity, while maintaining the number and identity of the mutations in any clone (54). For example, if mutations A and B were found together in the same sequenced clone, we retained their association throughout the procedure but randomly assigned the set to a different clone label. We calculated the specificity test statistic for each of 10,000 permutations of the clone labels. This procedure yields the expected distribution of the test statistic under the null hypothesis that the similarity among lines is independent of the antibiotic treatment or founding genotype. We then calculated an approximate *p*-value for rejecting this null hypothesis from the proportion of permutations in the expected distribution with a specificity statistic value greater than or equal to the observed value.

To quantify the specificity of genomic evolution with respect to genetic background, we performed an independent randomization test for each of the four antibiotics. Because these tests address the same null hypothesis, we combined the resulting *p*-values using Fisher’s method with 2*k* = 8 degrees of freedom, where *k* is the number of comparisons (29, 30). We provide the datasets and details of our statistical analyses in an R Notebook on GitHub (https://github.com/KyleCard/LTEE-ABR-mutant-sequencing).

## ACKNOWLEDGMENTS

We thank Chris Adami, Frances Downes, Joshua Franklin, Devin Lake, Rohan Maddamsetti, and Chris Waters for valuable discussions and advice. We acknowledge financial support from an HHMI Gilliam Fellowship (to K.J.C.); a Ralph Evans Award from the MSU Department of Microbiology and Molecular Genetics (to K.J.C.); a grant from the NSF (currently DEB-1951307 to R.E.L. and J.E.B.); a Hatch grant from the USDA (MICL02253 to R.E.L.); and the BEACON Center for the Study of Evolution in Action (NSF cooperative agreement DBI-0939454).

## Author contributions

K.J.C. and R.E.L. conceived of and designed the study; M.T. performed genomic DNA isolation and sequencing; K.J.C., J.L.G., J.E.B., and R.E.L. performed bioinformatics; K.J.C. and R.E.L. analyzed all data; K.J.C. wrote data-analysis scripts; K.J.C. and J.E.B. prepared figures; K.J.C., J.E.B., and R.E.L. wrote the paper. All authors approved the final version.

## Supplementary Table

**Table S1.** Bacterial strains sequenced in this study.

| Antibiotic* | LTEE parent<sup>†</sup> | Replicate | Strain |
| --- | --- | --- | --- |
| AMP | Ancestor | 1 | KJC108 |
| AMP | Ancestor | 2 | KJC109 |
| AMP | Ancestor | 3 | KJC110 |
| AMP | Ancestor | 4 | KJC111 |
| AMP | Ara-5 | 1 | KJC114 |
| AMP | Ara-5 | 2 | KJC122 |
| AMP | Ara-5 | 3 | KJC130 |
| AMP | Ara-6 | 1 | KJC115 |
| AMP | Ara-6 | 2 | KJC123 |
| AMP | Ara-6 | 3 | KJC131 |
| AMP | Ara+4 | 1 | KJC112 |
| AMP | Ara+4 | 2 | KJC120 |
| AMP | Ara+4 | 3 | KJC128 |
| AMP | Ara+5 | 1 | KJC113 |
| AMP | Ara+5 | 2 | KJC121 |
| AMP | Ara+5 | 3 | KJC129 |
| CRO | Ancestor | 1 | KJC212 |
| CRO | Ancestor | 2 | KJC213 |
| CRO | Ancestor | 3 | KJC214 |
| CRO | Ancestor | 4 | KJC215 |
| CRO | Ara-5 | 1 | KJC218 |
| CRO | Ara-5 | 2 | KJC226 |
| CRO | Ara-5 | 3 | KJC234 |
| CRO | Ara-6 | 1 | KJC219 |
| CRO | Ara-6 | 2 | KJC227 |
| CRO | Ara-6 | 3 | KJC235 |
| CRO | Ara+4 | 1 | KJC216 |
| CRO | Ara+4 | 2 | KJC224 |
| CRO | Ara+4 | 3 | KJC232 |
| CRO | Ara+5 | 1 | KJC217<sup>‡</sup> |
| CRO | Ara+5 | 2 | KJC225 |
| CRO | Ara+5 | 3 | KJC233 |
| CIP | Ancestor | 1 | KJC148 |
| CIP | Ancestor | 2 | KJC149 |
| CIP | Ancestor | 3 | KJC150 |
| CIP | Ancestor | 4 | KJC151 |
| CIP | Ara-5 | 1 | KJC154 |
| CIP | Ara-5 | 2 | KJC162 |
| CIP | Ara-5 | 3 | KJC186 |
| CIP | Ara-6 | 1 | KJC155 |
| CIP | Ara-6 | 2 | KJC163 |
| CIP | Ara-6 | 3 | KJC187 |
| CIP | Ara+4 | 1 | KJC152<sup>‡</sup> |
| CIP | Ara+4 | 2 | KJC160 |
| CIP | Ara+4 | 3 | KJC184<sup>‡</sup> |
| CIP | Ara+5 | 1 | KJC153 |
| CIP | Ara+5 | 2 | KJC161 |
| CIP | Ara+5 | 3 | KJC185 |
| TET | Ancestor | 1 | KJC60 |
| TET | Ancestor | 2 | KJC61 |
| TET | Ancestor | 3 | KJC62 |
| TET | Ancestor | 4 | KJC63 |
| TET | Ara-5 | 1 | KJC66<sup>§</sup> |
| TET | Ara-5 | 2 | KJC74 |
| TET | Ara-5 | 3 | KJC82 |
| TET | Ara-6 | 1 | KJC67 |
| TET | Ara-6 | 2 | KJC75 |
| TET | Ara-6 | 3 | KJC83 |
| TET | Ara+4 | 1 | KJC64 |
| TET | Ara+4 | 2 | KJC72 |
| TET | Ara+4 | 3 | KJC80 |
| TET | Ara+5 | 1 | KJC65<sup>§</sup> |
| TET | Ara+5 | 2 | KJC73 |
| TET | Ara+5 | 3 | KJC81 |

\*AMP, ampicillin; CRO, ceftriaxone; CIP, ciprofloxacin; TET, tetracycline.
<sup>†</sup>Ancestor, *E. coli* B strain REL606. All other parental strains are clones sampled at generation 50,000 from the indicated LTEE population.
<sup>‡</sup>Three strains (KJC152, KJC184, KJC217) are cross-contaminants, and they were discarded from all analyses.
<sup>§</sup>Two strains (KJC65, KJC66) have no identifiable mutations.

## References

1. S. Conway Morris, Life’s Solution: Inevitable Humans in a Lonely Universe (Cambridge University Press, 2003).

2. R. Woods, D. Schneider, C. L. Winkworth, M. A. Riley, R. E. Lenski, Tests of parallel molecular evolution in a long-term experiment with *Escherichia coli*. Proc. Natl. Acad. Sci. U.S.A. 103, 9107–9112 (2006).

3. R. E. Lenski, Convergence and divergence in a long-term experiment with bacteria. Am. Nat. 190, S57–S68 (2017).

4. G. Achaz, A. Rodriguez-Verdugo, B. S. Gaut, O. Tenaillon, “The reproducibility of adaptation in the light of experimental evolution with whole genome sequencing” in Ecological Genomics. Advances in Experimental Medicine and Biology, 781st Ed., C. R. Landry, N. Aubin-Horth, Eds. (Springer, Dordrecht, 2014), pp. 211–231.

5. Z. D. Blount, R. E. Lenski, J. B. Losos, Contingency and determinism in evolution: Replaying life’s tape. Science 362, eaam5979 (2018).

6. J. O’Neill, Tackling drug-resistant infections globally: Final report and recommendations (2016).

7. W. Wohlleben, Y. Mast, E. Stegmann, N. Ziemert, Antibiotic drug discovery. Microb. Biotechnol. 9, 541–548 (2016).

8. D. Nichol, et al., Steering evolution with sequential therapy to prevent the emergence of bacterial antibiotic resistance. PLoS Comput. Biol. 11, e1004493 (2015).

9. S. Baker, N. Thomson, F.-X. Weill, K. E. Holt, Genomic insights into the emergence and spread of antimicrobial-resistant bacterial pathogens. Science 360, 733–738 (2018).

10. D. Hughes, D. I. Andersson, Evolutionary trajectories to antibiotic resistance. Annu. Rev. Microbiol. 71, 579–596 (2017).

11. S. Iram, et al., Controlling the speed and trajectory of evolution with counterdiabatic driving. Nat. Phys., in press (2020).

12. E. Toprak, et al., Evolutionary paths to antibiotic resistance under dynamically sustained drug selection. Nat. Genet. 44, 101–105 (2012).

13. L. Imamovic, M. O. A. Sommer, Use of collateral sensitivity networks to design drug cycling protocols that avoid resistance development. Sci. Transl. Med. 5, 204ra132 (2013).

14. V. Lázár, et al., Bacterial evolution of antibiotic hypersensitivity. Mol. Syst. Biol. 9, 700 (2013).

15. M. Baym, et al., Spatiotemporal microbial evolution on antibiotic landscapes. Science 353, 1147–1151 (2016).

16. P. Yen, J. A. Papin, History of antibiotic adaptation influences microbial evolutionary dynamics during subsequent treatment. PLoS Biol. 15, e2001586 (2017).

17. K. J. Card, T. LaBar, J. B. Gomez, R. E. Lenski, Historical contingency in the evolution of antibiotic resistance after decades of relaxed selection. PLoS Biol. 17, e3000397 (2019).

18. J. Maltas, K. B. Wood, Pervasive and diverse collateral sensitivity profiles inform optimal strategies to limit antibiotic resistance. PLoS Biol. 17, e3000515 (2019).

19. D. Nichol, et al., Antibiotic collateral sensitivity is contingent on the repeatability of evolution. Nat. Commun. 10, 334 (2019).

20. T. Oz, et al., Strength of selection pressure is an important parameter contributing to the complexity of antibiotic resistance evolution. Mol. Biol. Evol. 31, 2387–2401 (2014).

21. G. Chevereau, et al., Quantifying the determinants of evolutionary dynamics leading to drug resistance. PLoS Biol. 13, e1002299 (2015).

22. L. Imamovic, et al., Drug-driven phenotypic convergence supports rational treatment strategies of chronic infections. Cell 172, 1–14 (2018).

23. R. E. Lenski, M. R. Rose, S. C. Simpson, S. C. Tadler, Long-term experimental evolution in *Escherichia coli*. I. Adaptation and divergence during 2,000 generations. Am. Nat. 138, 1315–1341 (1991).

24. O. Tenaillon, et al., Tempo and mode of genome evolution in a 50,000-generation experiment. Nature 536, 165–170 (2016).

25. O. Lamrabet, M. Martin, R. E. Lenski, D. Schneider, Changes in intrinsic antibiotic susceptibility during a long-term evolution experiment with *Escherichia coli*. mBio 10, e00189–19 (2019).

26. J. M. A. Blair, M. A. Webber, A. J. Baylay, D. O. Ogbolu, L. J. V. Piddock, Molecular mechanisms of antibiotic resistance. Nat. Rev. Microbiol. 13, 42–51 (2015).

27. I. M. Keseler, et al., The EcoCyc database: reflecting new knowledge about *Escherichia coli* K-12. Nucleic Acids Res. 45, D543–D550 (2017).

28. O. Tenaillon, et al., The molecular diversity of adaptive convergence. Science 335, 457–461 (2012).

29. R. A. Fisher, Statistical Methods for Research Workers, F. A. E. Crew, D. W. Cutler, Eds., 5th Ed. (Oliver and Boyd, 1934).

30. R. R. Sokal, F. J. Rohlf, Biometry: The Principles and Practices of Statistics in Biological Research, 3rd Ed. (W. H. Freeman and Company, 1994).

31. Z. D. Blount, C. Z. Borland, R. E. Lenski, Historical contingency and the evolution of a key innovation in an experimental population of *Escherichia coli*. Proc. Natl. Acad. Sci. U.S.A. 105, 7899–7906 (2008).

32. H. Aiba, T. Mizuno, Phosphorylation of a bacterial activator protein, OmpR, by a protein kinase, EnvZ, stimulates the transcription of the *ompF* and *ompC* genes in *Escherichia coli*. FEBS Lett. 261, 19–22 (1990).

33. S. Chakraborty, L. J. Kenney, A new role of OmpR in acid and osmotic stress in *Salmonella* and *E. coli*. Front. Microbiol. 9, 2656 (2018).

34. U. Choi, C.-R. Lee, Distinct roles of outer membrane porins in antibiotic resistance and membrane integrity in *Escherichia coli*. Front. Microbiol. 10, 953 (2019).

35. S. E. Meyer, S. Granett, J. U. Jung, M. R. Villarejo, Osmotic regulation of PhoE porin synthesis in *Escherichia coli*. J. Bacteriol. 172, 5501–5502 (1990).

36. K. Nishino, A. Yamaguchi, Role of histone-like protein H-NS in multidrug resistance of *Escherichia coli*. J. Bacteriol. 186, 1423–1429 (2004).

37. R. Gillet, B. Felden, Transfer RNA^Ala^ recognizes transfer-messenger RNA with specificity; a functional complex prior to entering the ribosome? EMBO J. 20, 2966–2976 (2001).

38. D. I. Andersson, D. Hughes, Antibiotic resistance and its cost: is it possible to reverse resistance? Nat. Rev. Microbiol. 8, 260–271 (2010).

39. The European Committee on Antimicrobial Susceptibility Testing, “Breakpoint tables for interpretations of MICs and zone diameters, version 10.0, 2020” (2020).

40. F. W. Studier, P. Daegelen, R. E. Lenski, S. Maslov, J. F. Kim, Understanding the differences between genome sequences of *Escherichia coli* B strains REL606 and BL21(DE3) and comparison of the *E. coli* B and K-12 Genomes. J. Mol. Biol. 394, 653–680 (2009).

41. A. P. Pugsley, J. P. Rosenbusch, OmpF porin synthesis in *Escherichia coli* strains B and K-12 carrying heterologous *ompB* and/or *ompF* loci. FEMS Microbiol. Lett. 16, 143–147 (1983).

42. D. Schneider, et al., Genomic comparisons among *Escherichia coli* strains B, K-12, and O157:H7 using IS elements as molecular markers. BMC Microbiol. 2, 18 (2002).

43. L. Sandegren, D. I. Andersson, Bacterial gene amplification: implications for the evolution of antibiotic resistance. Nat. Rev. Microbiol. 7, 578–588 (2009).

44. V. S. Cooper, A. F. Bennett, R. E. Lenski, Evolution of thermal dependence of growth rate of *Escherichia coli* populations during 20,000 generations in a constant environment. Evolution 55, 889–896 (2001).

45. A. M. Bolger, M. Lohse, B. Usadel, Trimmomatic: a flexible trimmer for Illumina sequence data. Bioinformatics 30, 2114–2120 (2014).

46. E. Afgan, et al., The Galaxy platform for accessible, reproducible and collaborative biomedical analyses: 2018 update. Nucleic Acids Res. 46, W537–W544 (2018).

47. J. E. Barrick, et al., Identifying structural variation in haploid microbial genomes from short-read resequencing data using *breseq*. BMC Genomics 15, 1039 (2014).

48. H. Jeong, et al., Genome sequences of *Escherichia coli* B strains REL606 and BL21(DE3). J. Mol. Biol. 394, 644–652 (2009).

49. C. Raeside, et al., Large chromosomal rearrangements during a long-term evolution experiment with *Escherichia coli*. mBio 5, e01377–14 (2014).

50. E. R. Moxon, P. B. Rainey, M. A. Nowak, R. E. Lenski, Adaptive evolution of highly mutable loci in pathogenic bacteria. Curr. Biol. 4, 24–33 (1994).

51. Z. D. Blount, J. E. Barrick, C. J. Davidson, R. E. Lenski, Genomic analysis of a key innovation in an experimental *Escherichia coli* population. Nature 489, 513–518 (2012).

52. X. Jiang, et al., Invertible promoters mediate bacterial phase variation, antibiotic resistance, and host adaptation in the gut. Science 363, 181–187 (2019).

53. Z. D. Blount, et al., Genomic and phenotypic evolution of *Escherichia coli* in a novel citrate-only resource environment. eLife 9, e55414 (2020).

54. D. E. Deatherage, J. L. Kepner, A. F. Bennett, R. E. Lenski, J. E. Barrick, Specificity of genome evolution in experimental populations of *Escherichia coli* evolved at different temperatures. Proc. Natl. Acad. Sci. U.S.A. 114, E1904–E1912 (2017).

